# Integrated environmental drivers and evolutionary history reveal conserved patterns of avian exposure to West Nile and Saint Louis encephalitis viruses

**DOI:** 10.1101/2025.10.22.683905

**Authors:** O. Giayetto, A.I. Quaglia, A.P. Mansilla, F.N. Nazar, A. Diaz

## Abstract

The increasing threat of emerging vector-borne diseases, affecting both humans and wildlife, necessitates a nuanced understanding of the complex interactions among vectors, hosts, and pathogens. Focusing on West Nile Virus (WNV) and Saint Louis encephalitis Virus (SLEV) activity in avian communities of central and northern Argentina, this study delves into the intricate dynamics of exposure risk employing phylogenetic mixed models. Contrary to expectations, ecological traits displayed negligible influence on exposure risk. Instead, temperature seasonality and altitude, emerged as crucial predictors. The phylogenetic signal indicated a conserved susceptibility along the avian evolutionary tree, with Furnariidae, Turdidae and Columbidae families exhibiting higher exposure risk. These findings challenge prevailing notions about the dominant role of ecological traits in mosquito-borne pathogens. The study underscores the need for a holistic understanding, emphasizing the intricate interplay of environmental, phylogenetic, and ecological elements in shaping avian exposure to WNV and SLEV, offering vital insights for future research and public health strategies.

## Introduction

Emerging vector borne diseases are a growing important health threat for humans, wildlife, and domesticated species, as a result from changes in the ecology of at least one of the actors involved in their maintenance in the ecosystem: vectors, hosts, or the pathogen itself (Daszak *et al*. 2000; Viglietta *et al*. 2021). When the pathogen is vector-borne, its transmission dynamic is shaped by how vector-host interactions are developed (Diaz *et al*. 2013b; Kilpatrick *et al*. 2006b; Vogels *et al*. 2019).

Identifying factors involved in the variation of host susceptibility at interspecific levels is crucial for understanding mechanisms of pathogen transmission and the suitability for pathogen amplification in the ecosystem (Yan *et al*. 2021). Resistance, tolerance, and exposure are essential factors within susceptibility for understanding the spread and impact of pathogens in host communities (Walsh 2019). On one hand, resistance and tolerance constitute important aspects of host competence (Burgan *et al*. 2019; Gervasi *et al*. 2015); as they are associated with host response to pathogen infection, influencing the host’s ability to support pathogen replication, while mitigating pathogen damage. On the other hand, the exposure risk plays a critical role prior to host infection, establishing the likelihood of host-pathogen interactions and pathogen vectorial transmission. These three factors, which determine the variation in species susceptibility, are influenced by the species’ particular characteristics. In essence, host’s traits can be related to the specieś ability to resist and tolerate while also impacting how vector-host interactions occur (Yan *et al*. 2021), shaping species infection risk as a result (Brinkerhoff *et al*. 2019; Huang *et al*. 2013; Valenzuela-Sánchez *et al*. 2021; Yan *et al*. 2017).

Host attributes are influential factors in predicting infection risk to vector-borne pathogens, providing information about how the ecological processes are structured and offering insights with direct implications for human health (Daszak *et al*. 2000; Habarugira *et al*. 2020). Studies have identified that body size (Figuerola *et al*. 2008; Gutiérrez-López *et al*. 2019; Lutz *et al*. 2015; Skinner *et al*. 2021), migratory status (Chevalier *et al*. 2009; Figuerola *et al*. 2008), nest characteristics (González *et al*. 2014; Lutz *et al*. 2015) and reproductive and developmental traits (Ganser *et al*. 2020; Walsh 2019) are important predictors of host infection risk to vector-borne pathogens. This suggests that the distribution of infection risk and species life traits strategies follow similar trajectories, they can directly affect the way and the extent to which host-vector interaction occurs by differentially exposing hosts to mosquito bites.

Host infection risk could be phylogenetically conserved when assessing the influence of host traits. Confounding effects in identifying important traits could arise from disregarding the phylogenetic relationships between hosts (Barrow *et al*. 2019; Gupta *et al*. 2020). For instance, this conservation of infection risk could stem from phylogenetically conserved traits among hosts, constraints imposed by host phylogeny on the pathogen’s host range, or even the co-evolutionary history between vector and host traits (Lutz *et al*. 2015; Valenzuela-Sánchez *et al*. 2021). Combining life traits and phylogeny of host species in the analysis of vector-borne pathogens infection risk have been encouraged to deeply understand the mechanisms beneath interspecific variation in infection risk (Barrow *et al*. 2019; Gupta *et al*. 2020; Pigeault *et al*. 2022).

Moreover, environmental characteristics at regional and local scale have been related to pathogen transmission as they can impact on the behaviour, distribution, and survival of vector and host directly, affecting the pathogen prevalence in the host community (Giesen *et al*. 2023; González *et al*. 2014; Levine *et al*. 2016). Factors such as temperature, humidity and precipitation are associated with the risk of transmission of vector-borne pathogens, as these local features mainly influence vector development, reproduction and survival (Giesen *et al*. 2023; Tabachnick 2010). On the other hand, landscape features influence where and when competent hosts and vectors interact, configuring suitable transmission hotspots in specific environmental scenarios (Hess *et al*. 2018).

Particularly, West Nile Virus (WNV) and Saint Louis Encephalitis Virus (SLEV) are two emerging arboviruses belonging to the family Flaviviridae. Both are causative agents of encephalomyelitis and are considered of global health concern (Guth *et al*. 2020), affecting not only humans but also wildlife and cattle (LaDeau *et al*. 2007; Ong *et al*. 2021). West Nile virus and SLEV are maintained within a complex multi-vector, multi-host transmission network shaped by intricate ecological interactions and modulated by spatial and seasonal fluctuations in viral transmission (Diaz *et al*. 2013b). Both viruses share an ecological resemblance in their maintenance network being predominantly maintained by *Culex* mosquitoes and a wide range of avian species (mainly Passeriformes and Columbiformes) (Ciota 2017; Diaz *et al*. 2013a; Giayetto *et al*. 2021; Reisen 2003; Rochlin *et al*. 2019).

Like most of the flavivirus, WNV and SLEV produce short term viremias in avian hosts that persist detectable for no more than 10 days (Reisen 2003, 2013), and only WNV has been shown to be pathogenic for some bird species in the Palearctic and Nearctic (Diaz *et al*. 2011). For the southern region of South America, partial knowledge exists regarding the vectors and hosts involved in the SLEV and WNV maintenance, and information about their vector’s feeding preferences is limited. Both viruses can be vectorized at least by four species of *Culex* mosquitoes (*Culex quinquefasciatus* Say, *Culex pipiens*, *Culex interfor* and *Culex saltanensis*) (Diaz *et al*. 2013a; Giayetto *et al*. 2021), including columbiform and passeriform birds as feeding resources (Cardo & Vezzani 2023; Cardo *et al*. 2023). Regarding hosts, Eared doves (*Zenaida auriculata*), Picui ground-doves (*Columbina picui*) and House sparrows (*Passer domesticus*) species have developed infectious viremias for both viruses (Diaz *et al*. 2018, 2011). Although WNV and SLEV are two arboviruses widely studied few studies have focused on identifying risk factors for their potential hosts. In the case of WNV, previous research has highlighted that migratory movements, nesting strategies, and body size are traits that increase risk within the Afro-Palearctic maintenance and amplification system (Chevalier *et al*. 2009; Figuerola *et al*. 2007). However, similar approaches have not yet been conducted for SLEV. Environmental and landscape features such as rural areas, wetlands, and small urban zones (Lockaby *et al*. 2016; Mansilla *et al*. 2022; Reisen 2013), alongside increments in temperature and humidity during warmest months (García-Carrasco *et al*. 2024; Hess *et al*. 2018) have consistently been associated with a higher transmission risk for both WNV and SLEV.

In this context, where behavioural, ecological, environmental, and evolutionary factors influence viral transmission, we aimed to elucidate their relative roles and assess whether they increase or reduce the risk of host infection by these vector-borne viruses. For this study, serosurveys for WNV and SLEV informed infection history in wild birds communities encompassed among a broad array of biogeographic units and landscapes in central and northern Argentina. We used phylogenetic Bayesian mixed models to study whether host life history strategy, ecological features or environmental conditions associate hosts infection risk to WNV and SLEV while considering the potential effects of host phylogenetic relationships.

## Methods

### Seroprevalence data

A long term and regional cross-sectional serological study was conducted to detect the exposure of free-ranging birds to WNV and/or SLEV gathered in published (Diaz *et al*. 2008; Mansilla *et al*. 2022) and unpublished data between 2004 and 2019. The serosurveys encompassed 31 sampling events in 19 locations along subtropical to temperate climates and a wide range of pristine to human-modified environments on 7 biogeographic units in north and central Argentina (latitude −26.82° and −36.64°) (Figure 1; Table 1).

**Figure 1.**
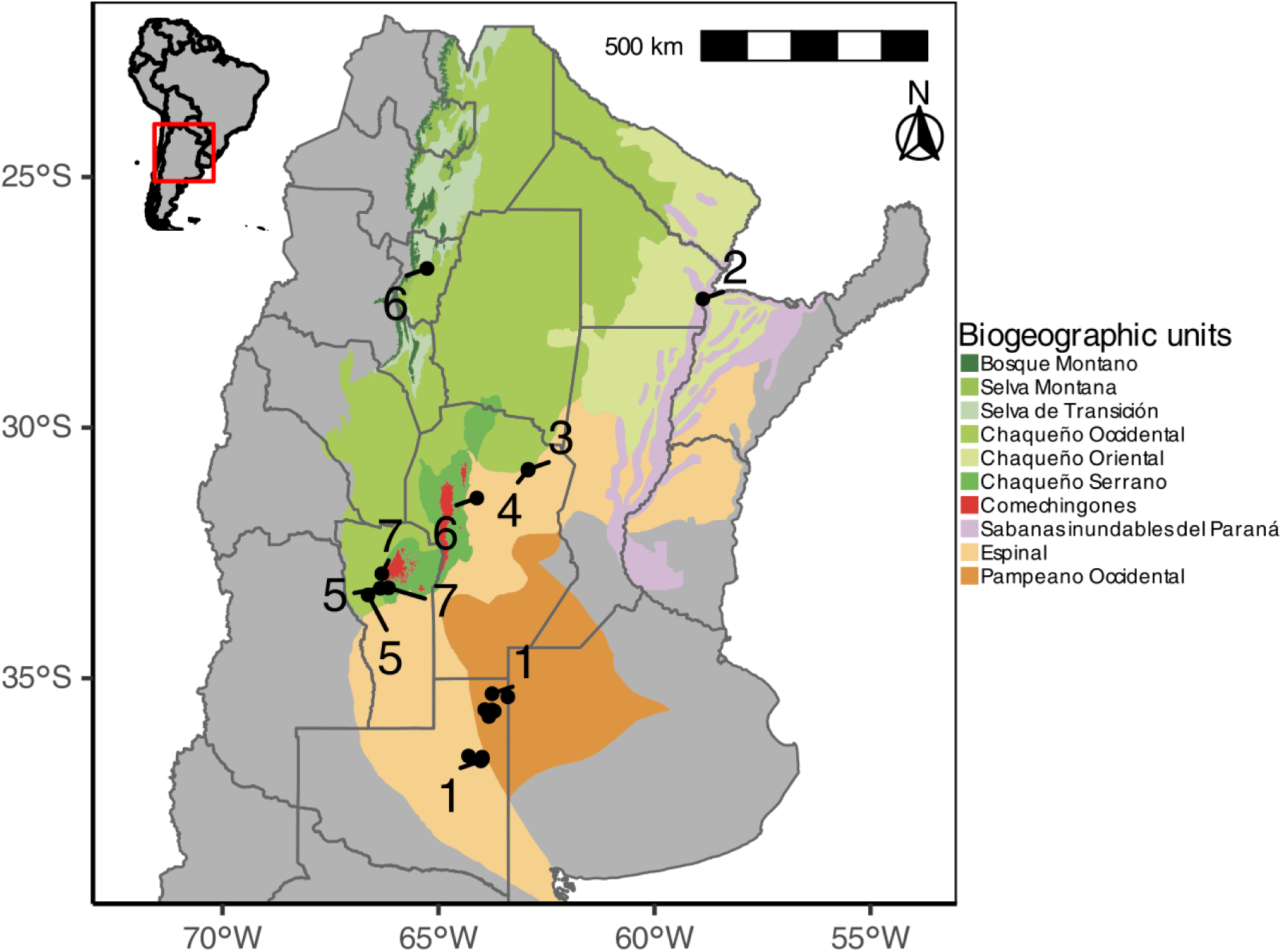
Map of sampled Argentinean biogeographic units. Each dot represents a sampling event. Sampling events were clustered to represent 7 transmission environments. Cluster characteristics are summarized in Table 1. The inset map shows South America, with central and northern region of Argentina highlighted in the red box.

**Table 1.** Summarized information of the transmission environments after cluster analysis results. The table displays the biogeographic units and the number of sampling events included in the cluster, along with the general climatic characteristics, the number of individuals and the species sampled within the transmission environment.

| Cluster | Biogeographic unit | Climate | Habitat | Sampling events | Sampled birds | Sampled species |
| --- | --- | --- | --- | --- | --- | --- |
| 1 | Pampeano<br>Occidental-Espinal | Temperate, no dry<br>season, hot summer | Agroecosystem | 10 | 809 | 27 |
| 2 | Chaqueño Oriental-Sabanas<br>inundables del<br>Paraná | Temperate, no dry<br>season, hot summer | Urban | 2 | 122 | 17 |
| 3 | Espinal | Temperate, no dry<br>season, hot summer | Wetland | 3 | 385 | 34 |
| 4 | Espinal | Temperate, no dry<br>season, hot summer | Dry forest | 7 | 1008 | 41 |
| 5 | Chaqueño<br>Occidental | Arid, steppe, cold | Dry forest | 2 | 122 | 14 |
| 6 | Espinal-Chaqueño<br>Occidental | Temperate, dry winter, hot<br>summer | Urban | 5 | 692 | 36 |
| 7 | Chaqueño Serrano | Arid, steppe, cold | Highland dry forest | 2 | 72 | 13 |

The presence of neutralizing antibodies in avian sera was used as indicative of a previous exposure to WNV and/or SLEV (*i.e.,* survival). Birds exposed to WNV and SLEV develop an acute infection with viremias detectable during a few days and long-lasting antibodies detectable for two years (Gibbs *et al*. 2005; McKee *et al*. 2015), then, the presence of neutralizing antibodies is suitable to evaluate the risk of exposure in birds in cross-sectional surveys.

Birds’ sampling procedure was performed as described elsewhere (Diaz *et al*. 2016). Briefly, birds were captured using mist nets, manually restrained, ringed and bleed by jugular venipuncture. Blood was allowed for clotting at room temperature and then centrifuged for serum collection. Sera samples were stored at −20°C The presence of WNV and SLEV-reactive antibodies in sera samples were analysed by a plaque-reduction neutralization test with a serum dilution threshold that neutralised 80% of the virus. This is the gold standard serological test in arbovirology and a suitable protocol to evidence birds that were exposed to both viruses (Gilbert *et al*. 2013). Cross-reactivity with other flavivirus was taken into consideration when performing the neutralization test (Tauro *et al*. 2012).

### Ecological and life-history traits data

We compiled a dataset of ecological and life-history traits that were potentially relevant to explain WNV and SLEV birds’ exposure risk (Figuerola *et al*. 2008; Ganser *et al*. 2020; Lutz *et al*. 2015).

#### Life history traits

Species were ranked on the pace of life continuum by means of a principal component analysis (PCA) performed with 9 traits related to reproduction (egg mass, clutch size and broods per year), size (wing length, tarsus length and body mass) and development (incubation period, generational length and fledgling period). Then, the pace of life strategy was synthesized by the first PCA axis (54.8% of inertia), hereafter mentioned as paces of life. This axis, and thus the new synthetic variable, represents the spectrum between fast (negative values) and slow (positive values) strategies (For further details see Supplementary material SM1).

#### Ecological traits

The following ecological traits were selected: nest height, nest type and migratory status. Nest height was classified as ground, understory (nests predominantly found above ground and below 3m) or canopy/subcanopy level (nests predominantly found above 3m). Nest type was classified as open cup, closed cup or cavity (Cooke *et al*. 2020; Lutz *et al*. 2015). Migratory status was classified as sedentary, partially migratory, and migratory (Tobias *et al*. 2022).

Species’ nest and eggs characteristics were extracted from De La Peña, 2023. Species data for body mass, clutch size, wing length, tarsus length, and migratory status was extracted from Tobias *et al*. 2022. The paucity of trait data for the assemblage of bird analysed here was filled with additional sources (Cooke *et al*. 2019; De Mársico *et al*. 2010; Mason 1986; Murton *et al*. 1974; Stott 1948; Wiley 1988).

### Environmental scenarios of viral transmission

Environmental information was considered to model the risk of infection regarding the nested nature of the SLEV and WNV transmission among the landscapes and biogeographic units (Diaz *et al*. 2013b; Giesen *et al*. 2023), in addition to include the spatial and temporal extent of sampling events (Figure 1; Table 1). A principal coordinates component analysis (PCoA) was used to characterise the environmental scenarios of viral transmission (ESVT). Two products were exploited from the PCoA: 1-ordination of the sampling events in clusters resembling ESVT which served as group-level intercept variable to model the risk of infection; and 2-synthetic variables summarising the complexity of the environmental features at local scale that could have an impact on the infection risk (*i.e.,* population-level slopes). To perform the PCoA the data included nine variables accounting monthly, seasonal and annual bioclimatic features (10 arc-minutes resolution; https://www.worldclim.org/), altitude, the identity of biogeographic unit (Oyarzabal *et al*. 2018) and the land use (urban and agroecosystems) for a given sampling event. Eight PCo axes contained the 80% of the variability, which were selected to identify the ESVT by means of *K*-means cluster analysis. Seven ESVT were the optimal number of clusters that minimize the total intra-cluster variation by the *elbow* method (Borcard *et al*. 2018). The first and second PCo axes represented the 40.3% and the 28.4% of the environmental variability, respectively (For further details see Supplementary material SM1). Inspection of the loadings showed that PCo_1_ was mainly correlated with the increase in the annual range of temperature (r = 0.94), hereafter accounting for the temperature seasonality of the sites, while PCo_2_ was negatively correlated with the altitude (r = - 0.88).

### Analysis of the ecological determinants of the infection risk

The effects of host traits and the environmental scenarios on bird infection risk for WNV and SLEV were analyzed with bayesian phylogenetic generalized linear multilevel models (B-PGLMMs). B-PGLMMs are useful to account for various sources of uncertainty and correlation in the data while accounting for species evolutionary history (Fountain-Jones *et al*. 2018; Pigeault *et al*. 2022). The WNV and SLEV serological status of the 49 species with more than 10 sampled individuals (n= 3,210) constituted the binomial response variables that informed on the risk of infection in the B-PGLMMs (Kernbach *et al*. 2021).

To infer the overall contribution of hosts traits and environmental scenarios as predictors, four models were compared for each virus (Table 2):

1- a Null model included group-level effects (intercepts) accounting for variation in the infection risk related to the ESVT, the phylogenetic constraints given the evolutionary relationship between species and intraspecific variation among sampling events;
2- an Ecological traits model considering only the host traits (migration status, the nest type and height, clutch size and the pace of life) as population-level slopes plus the group-level effects of the Null model;
3- an Environmental model considering only the environmental features at local scale (temperature seasonality and altitude) as population-level slopes plus the group-level effects of the Null model; and
4- a Full model considering the additive effects of host traits and the environmental features at local scale, plus the group-level effects of the Null model. The performance of the models was assayed with the approximate leave-one-out cross-validation information criterion based on the posterior likelihoods (LOOIC) (Vehtari et al. 2017).

**Table 2.** Comparison of four Bayesian phylogenetic generalized linear multilevel models for each virus. All models included the group-level effects (Species, transmission environment and phylogeny). Null model was fitted with no population level variables. The ecological trait model was fitted using nest height, nest type, pace of life, migratory status, and clutch size as population level variables. The environmental model only included temperature seasonality and altitude as population level variables. The full model included both ecological traits and environmental variables. Model fit was assessed using leave-one-out cross-validation and the difference between each model and the best fitted model is shown as ΔLOOIC with their standard error (SE).

| | ΔLOOIC (SE) | |
| --- | --- | --- |
|  | St. Louis Encephalitis virus | West Nile virus |
| Environment | 0(0) | 0(0) |
| Full | 1.6(3.4) | 1.1(3.4) |
| Ecological traits | 40.6(10.8) | 9.8(6) |
| Null | 40.8(10.1) | 9(5.1) |

No collinearity was detected between host traits and environmental predictors (Pearson correlation coefficients r ≤ |0.8|). Data exploration showed that the relationship between the infection risk and temperature seasonality was inverted U-shaped and a quadratic term was added. Continuous variables were standardized (mean=0; SD=1). The phylogenetic relationship between species was informed by means of a consensus tree (pack *phytools,* (Revell 2012)) based on 1,000 trees from BirdTree.org (Jetz *et al*. 2012), using the backbone tree (Hackett *et al*. 2008) in accordance with similar risk analysis (Barrow *et al*. 2019; Pigeault *et al*. 2022).

Default priors were set up in the B-PGLMMs and ran four chains with 20,000 iterations (10,000 iterations burn-in, thinned 10; package *brms* (Bürkner 2017)). Convergence of the four chains to a common stationary posterior distribution for the population and group-level parameters was assayed. In addition, the independence of the sampled values of the posterior distributions and the goodness of fit were checked using randomized quantile residuals (Inchausti 2022).

The potential confounding effect of the species trait-environmental relationship (Dray & Legendre 2008) was considered by exploring the distribution of bird species, families and traits along environmental clusters. Poisson generalized regression models (GLM) were fitted and the type II Wald’s *X^2^* analysis of deviance performed (*α*=0.05) to inspect the variation in the number of species and families surveyed between ESTVs. The homogeneity in the distribution of species traits between ESVT was tested with the fourth-corner analysis as implemented in Borcard et al. (2018) (9999 permutations and false discovery rate adjustment).

Environmental and host traits effects were inferred from the posterior distributions for each predictor in the Full models (median and 95% highest density credible interval, CI; packages *tidybayes* (Kay 2024). Predictors with CI not including zero were considered to have relevant influence on the risk of infection. The overall contribution of the group level effects to the infection risk was evaluated by variance partitioning (Barrow *et al*. 2019). The phylogenetic signal (λ) in the risk of infection was computed as the proportion of the total variance attributed to the phylogenetic distance between hosts (*Barrow et al. 2019; Pigeault et al. 2022*; Bürkner (https://cran.r-project.org/web/packages/brms/vignettes/brms_phylogenetics.html). GLMs were fitted with *glmmTMB* package (Brooks *et al*. 2017) and the fourth-corner analysis implemented with the *ape* packages (Paradis & Schliep 2019). All the analyses were performed in R (R Core Team 2021) and figures deployed with ggplot2 package (Wickham 2011).

## Results

A total of 3210 tested birds in 49 species, 16 families and 6 orders were included in our analysis. WNV seroprevalence accounted for 2.2% [95% CI 1.8 - 2.8]) with 61.22% (30/49) of the species exposed (Fig. 2). On the other hand, 6.3% [5.6 - 7.3]) of the tested birds were positive to SLEV with 77.55% (38/49) of the species exposed (Figure 2). Only 1.6% of the avian sera tested showed heterotypic immunological reaction indicating birds had been exposed to both viruses. Species that were exposed to WNV most frequently were *Drymornis bridgesii (Furnariidae)* (35.7%; [95% CI 16 - 61]), *Leptotila verreauxi* (18.8%; [95% CI 7 - 43), *Tarphonomus certhioides (Furnariidae)* (16.6%; [95% CI 5 - 48]), *Furnarius rufus (Furnariidae)* (13.9%; [95% CI 10 - 19]) and *Turdus rufiventris (Turdidae)* (12.0%; [95% CI 5 - 24]). Species most exposed to SLEV were *D. bridgesii* (42.9%; [95% CI 21 - 67]), *L. verreauxi* (31.2%; [95% CI 14 - 55]), *T. rufiventris* (22.0%; [95% CI 13 - 35]) and P*seudoseisura lophotes (Furnariidae)* (20.0 %; [95% CI 5 - 51]). As indicative of the compartmentalization in virus-host interactions, the family Bucconidae was only exposed to WNV (*Nystalus maculatus* 12.5%; [95% CI 0 - 30.5]) and 4 out 16 families surveyed were only exposed to SLEV (Mimidae, Passerellidae, Picidae and Scolopacidae).

**Figure 2.**
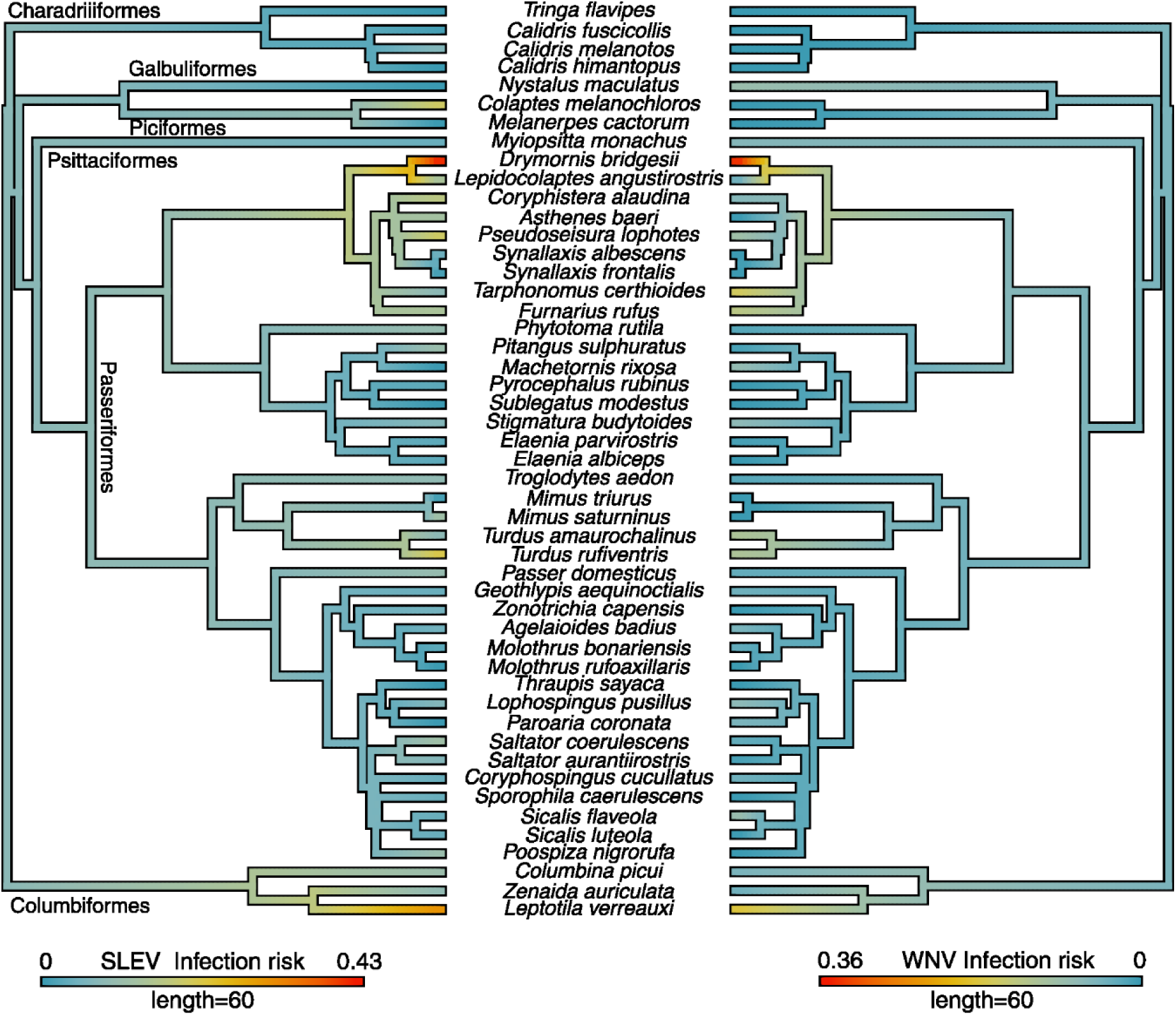
West Nile virus (WNV, right) and St. Louis Encephalitis virus (SLEV, left) prevalence in birds across the avian phylogeny, mapping the proportion of infected individuals for each well-sampled host species (≥ 10 individuals). A consensus tree was generated based on 1000 trees from BirdTree.org.

The variations in the amount of bird species, families and species life-traits surveyed were reasonable at the biogeographic scale analysed here. Accordingly, the number of species in the ESVT of the southern zone was up to three times lower than that observed in the ESVT of the central-northern zone of the study area (*X*^2^=58.37; df=6; p<0.001), with no variation at family level (*X*^2^=9.80; df=6; p=0.133). In addition, the distribution of the bird traits selected in this study were similarly distributed in the ESVT (For further details see Supplementary material SM2).

In regards to the overall contribution of hots traits and the context given by environmental scenarios where SLEV and WNV exposure occurred there were major differences informed by the performance of the tested models. Model comparison by LOOIC showed that models including only environmental predictors were the best fitted for both WNV and SLEV and performed slightly better than the full models with both environmental and ecological traits (ΔLOOIC < 1.5 for SLEV and WNV). Models that included only ecological traits fitted substantially worse than models with only environmental variables (ΔLOOIC > 39 for SLEV; ΔLOOIC > 8 for WNV). The ecological traits models performed similar to the null model for both viruses, indicating that the relevance of ecological traits evaluated were negligible (Table 2). Nevertheless, the following results were based on the full model in order to show the relative importance of each of the evaluated components (behavioural, ecological, environmental, and evolutionary).

The bird traits for global level effects included (migration status, the nest type and height, clutch size and the pace of life) showed negligible effect in the exposure risk for SLEV and WNV, as all 95% CI encompassed 0 (Figure 3). As indicated by the fourth corner analysis, the lack of effect of the tested traits could not be obscured by the distribution of traits across the ESVT.

**Figure 3.**
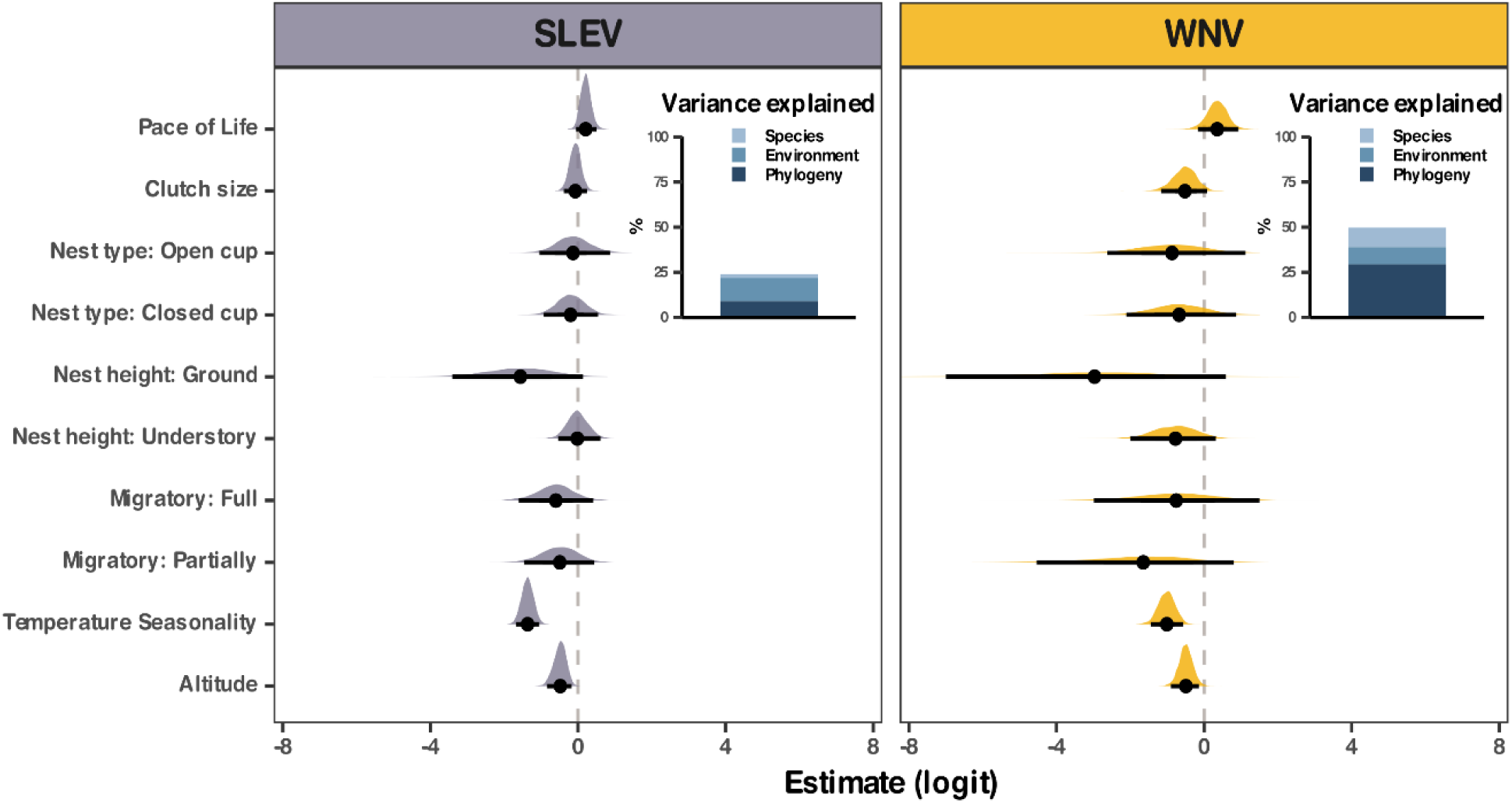
Posterior means estimates and credible intervals of population level effects on exposure risk to West Nile virus (WNV, right) and St. Louis Encephalitis virus (SLEV, left) for the full Bayesian phylogenetic generalized linear multilevel model (including environmental and ecological traits as response variables). For categorical predictors, the effects shown are relative to the reference categories: Nest type (Cavity), Nest height (Canopy/Subcanopy) and Migration (Sedentary). Dots denotate the posterior means and lines represent 50, 75 and 95% credible intervals. Shaded area shows posterior distribution of each population level effect. Panels illustrate the total variance attributed to group level effects: Species (lightblue), Environment (as transmission environments, darkblue) and phylogeny (brown).

The environmental global level effects were related with the reduction of the avian exposure risk to SLEV and WNV (Fig. 3). The quadratic term for temperature seasonality was an important predictor of birds’ exposure risk to WNV (logit scale: −1.02; [95% CI −1.48 - (−0.59)]) and SLEV (−1.36; [95% CI −1.68 – (−1.05)]). Birds’ exposure risk to both viruses was lower at the extreme’s values of temperature seasonality, meaning that the exposure risk is negatively affected by both extreme variations between seasons and minimal differences in temperature across the year (Fig. 4). Altitude was also an important predictor for SLEV (logit scale: −0.49; [95% CI −0.85 - (−0.19)]) and WNV (logit scale: −0.49; [95% CI −0.88 - (−0.12)]), showing that the exposure risk decreases with increasing altitude. However, the effect of altitude on exposure risk was more uncertain for WNV than for SLEV (Fig. 3 and 4). The environmental and evolutionary context constrained the prevalence of exposure to SLEV and WNV differently. The inherent hierarchy of the group level effects informed these findings (Fig. 3: inset barplot). The proportion of variance attributed to the group-level effects was 23.85% for SLEV and 49.35% for WNV. Out of total, the proportion of variability attributed to the ESVTs was 12.59% [95% CI 1.54 - 41.2]) for SLEV and 9.46 [95% CI 1.19 - 32.5]) for WNV, indicating that prevalence of exposure to each virus across different transmission environments had similar magnitude, and a similar trend between viruses within the same environment. The posterior median of the prevalence to SLEV was higher in the Espinal - Pampeano Occidental region (ESTV 1) and lower in the Chaqueño Occidental region (ESTV 5). The posterior mean of the prevalence to WNV was higher in the Chaqueño Oriental, Espinal and Pampeano regions (ESTVs 1, 2 and 3), and lower in the urban localities in the Espinal - Chaqueño Occidental region (ESTV 6) (For further details see Supplementary material SM2). The variance attributed to intraspecific variation in the prevalence of exposure was lower for SLEV than for WNV (2.26%, 95% CI [0–9.91] vs. 10.89%, 95% CI [2.0–37.1], respectively). This indicates that the tested species were consistently exposed to SLEV across sampling events in the ESTV, despite being sampled in different ESTVs, whereas their exposure to WNV showed greater variability. The phylogenetic signal in the exposure risk for both viruses was lower to SLEV compared to WNV, indicating that exposure risk is more conserved, to some extent, along avian evolutionary tree for WNV than SLEV. The variance attributed to the phylogenetic constraints was 9% [95% CI 2 - 26] for SLEV and 29% [95%CI 0.21 - 65] for WNV.

**Figure 4.**
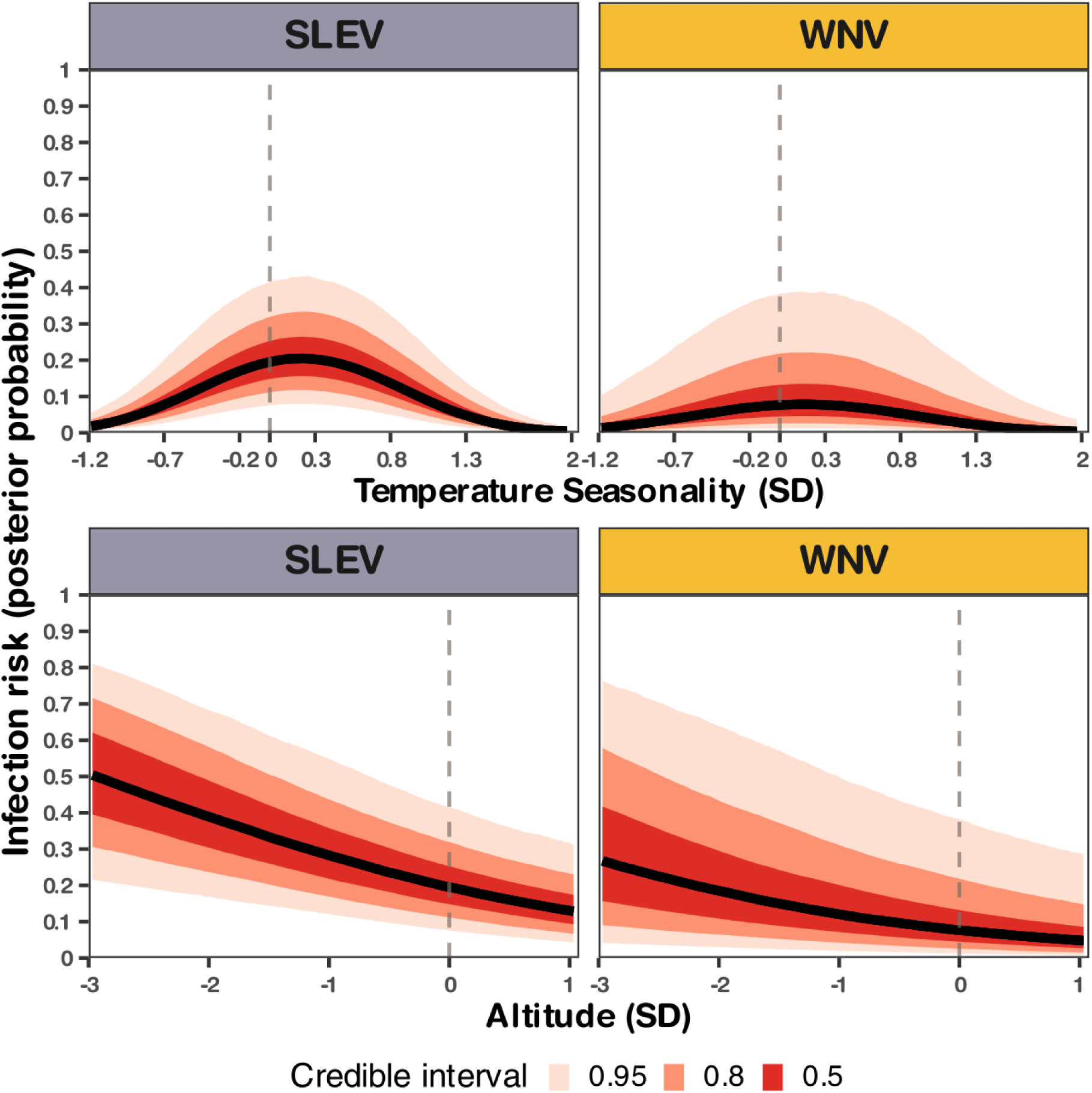
Conditional mean plots of the full Bayesian phylogenetic generalized linear multilevel model including both population- and group-level effects, showing the predicted relation between the infection risk in relation to (standardized) temperature seasonality (top) and altitude (bottom) for West Nile virus (right) and St. Louis Encephalitis virus (left). The colours represent credible intervals of 50, 80 and 95%.

## Discussion

In this study, we evaluate how species traits, their phylogenetic relationships, and the environmental conditions where viral transmission occurs affect the risk of exposure to SLEV and WNV. We found that ecological and life history traits did not influence over birds’ exposure risk to WNV and SLEV, but instead, environmental characteristics were predominantly more relevant. Besides, the variance explained by the phylogenetic signal was a particularly important factor in predicting the exposure risk to WNV, indicative of a conserved susceptibility along the avian evolutionary tree.

Bird exposure to WNV and SLEV was not significantly influenced by ecological or life history traits. Specifically, traits such as migratory status, clutch size, nesting characteristics and the pace of life did not appear to affect exposure risk in our study. These findings contrast with our expectations. We hypothesized that full migrant birds would be at higher risk, given their overlap with periods of peak vector activity and their use of diverse stopover sites, as reported in other geographic systems (Jourdain *et al*. 2007; Rappole *et al*. 2000; Swetnam *et al*. 2018). Likewise, we expected that nest characteristics would modulate exposure by placing birds at different heights—potentially aligning them with mosquito flight zones—or by offering greater or lesser structural protection during periods of limited mobility for both adults and fledglings. We also anticipated that larger-bodied birds would show increased exposure due to stronger olfactory and visual cues for mosquito detection. These hypotheses were based on a substantial body of literature, showing that avian ecological and life history traits are predictive of infection risk in mosquito-borne pathogen systems. For instance, nest height and structure have also been shown to influence mosquito access and, consequently, host exposure (Balenghien *et al*. 2006; Marra *et al*. 2004; Takken & Verhulst 2013). Migratory behaviour, clutch size, diet type, and pace of life have all been linked to variation in host exposure and infection in systems involving hemoparasites and arboviruses (Ganser *et al*. 2020; González *et al*. 2014; Pigeault *et al*. 2022; Skinner *et al*. 2021; Walsh 2019). Moreover, negative correlation between the pace of life and host competence, was observed in WNV and other arboviruses (Downs *et al*. 2019; Huang *et al*. 2013) and for others pathogen-hosts systems (Albery & Becker 2021; Johnson *et al*. 2012b). In the Afro-Palearctic system, migratory birds arriving from Africa were found to be more exposed to WNV than resident birds in Spain, possibly due to the differential viral activity along their routes (Chevalier *et al*. 2009; Durand *et al*. 2017; Figuerola *et al*. 2008). However, we observed a slight positive relationship between exposure risk and pace of life, possibly due to the high association with body mass. If this pattern becomes more significant in future studies, it may help to establish links between the exposure risk of potential host species and their ability to amplify these viruses. Overall, the absence of significant associations observed in our study suggests that the mechanisms described above may not operate in the same way at the southernmost distribution range of these viruses in the Neotropics.

In our models, most of the explained variance was associated with phylogeny, matching or even exceeding the variance attributable to environmental factors. This strong phylogenetic signal in exposure risk was primarily driven by the families Furnariidae and Columbidae. Our findings are consistent with Tolsá et al., 2018 (Tolsá *et al*. 2018), who identified Columbiformes and Passeriformes as the orders phylogenetically most exposed to WNV worldwide. Notably, the Furnariidae family is restricted to the Neotropical region, whereas the *Flavivirus* genus (including SLEV and WNV) originated in Africa (Pettersson & Fiz-Palacios 2014). This suggests that the conservation of exposure risk in these multi-host, multi-vector viruses is more likely driven by vector–host co-adaptation mediated by feeding behaviour and habitat use than by shared evolutionary history with the viruses themselves (Lyimo & Ferguson 2009; Takken & Verhulst 2013).

It is well established that environmental conditions shift vector borne pathogens transmission by affecting virus-vector-host interactions (Giesen *et al*. 2023). Particularly for WNV and SLEV, several factors such as land use, temperature, humidity, precipitation, and altitude have been mainly associated with virus prevalence in birds and vector abundance (Mansilla *et al*. 2022; Marcantonio *et al*. 2015; Shaman *et al*. 2002). We found that the exposure risk of birds to WNV and SLEV was better explained by their transmission environment than by their life traits, with temperature seasonality and altitude being particularly influential. It was observed that, for both maximum and minimum temperatures variation, viral activity was lower at the extreme ranges compared to intermediate temperatures, which aligns with the seasonal temperature patterns in the study area. On the other hand, SLEV and WNV exposure risk were negatively correlated with altitude. In the northern regions of Argentina, temperature seasonality is less pronounced and reaches its maximum towards the south. Therefore, we could expect that the exposure risk of birds to WNV and SLEV would be highest in the central region of Argentina, which is characterized by mild temperate conditions and low altitudes. Some of the variation in exposure risk was associated with the ESVTs, where environments like agroecosystems in the Pampeano Occidental region or urban environments of the Chaqueño oriental region were slightly more suitable for SLEV and WNV activity. This pattern partially aligns with previous literature, which describes how agricultural and urban areas often facilitate the transmission of these viruses (Giesen *et al*. 2023; Hess *et al*. 2018; Kovach & Kilpatrick 2024; Mansilla *et al*. 2022). However, we did not observe this increased risk in the ESVTs associated with another environment highly linked in the literature to SLEV and WNV activity (i.e. wetlands and subtropical forest) (Johnson *et al*. 2012a; Sánchez-Gómez *et al*. 2017). The central region of Argentina (Pampeano Occidental region) is one of the most disturbed areas where landscape is dominated by agriculture and urban environments. Such environments harbouring low biodiversity indexes which can promote viral activity from the dilution effect perspective (Keesing *et al*. 2006). Alternatively, the exposure risk for WNV varied among species depending on the ESVT in which it was assessed. This could be due to vector-host interactions being influenced by the bioclimatic conditions of the environment, either through changes in vector feeding patterns shared among ESVTs (Abella-Medrano *et al*. 2018; Kilpatrick *et al*. 2006a), or through inherent differences in vector communities across ESVTs (Cardo *et al*. 2012; Stein *et al*. 2016).Regardless of the underlying reason, the variation in a species’ exposure risk relative to its ESVT can directly influence the maintenance network of these viruses, assigning greater or lesser relevance to potential host species. Overall, these results highlight the importance of considering bioenvironmental conditions when assessing the exposure risk to SLEV and WNV in any studied region.

Our study summarizes and presents an extensive collection of serological surveys conducted on birds from different environments in the southernmost distribution zone of these viruses. Furthermore, based on this compilation of serological data, we were able to look into the predominant causes that shapes birds’ exposure risk to SLEV and WNV, despite their relative low prevalence in nature. However, these results should be interpreted with caution, as sampling structure and capture procedures may introduce bias. These factors could distort the diversity of analysed transmission environments and the structure of the trait’s matrix, even though traits are evenly represented across environments. Our species pool may not be diverse enough, possibly due to our capture protocol using mist nets (Tattoni & LaBarbera 2022). Most of the sampled species were underrepresented, from 132 species captured only 49 had at least 10 individuals sampled, but this fact could be recurrent in arbovirus risk studies of similar magnitudes (Durand *et al*. 2017). Taking this into account, we can conclude that environment and phylogeny are significant determinants in the processes involved in the transmission and maintenance of WNV and SLEV in the studied region, achieving higher exposure risk in species from Furnariidae, and Columbidae families in the central regions of the country where temperature seasonality is more pronounced. These findings highlight the need for studies capable of assessing how the interface between the environment and host selection by mosquito vectors unfolds in the context of viral transmission, guiding future studies and prevention strategies to avoid potential escalations in both human and animal cases.

## Supporting information

Codes

## Acknowledgments

We are grateful to M.D. Beranek, A. Visintin, M. Laurito, M. Stein, G. Oria, J. Rosas, MJ Dantur Juri, R. Lobos, E. Seiler, S. Flores and Batallán, PG for their technical assistance during sample collection and serological determinations. This project was supported by grants PICT 2018-01172, SECYT Consolidar UNC and PUE 2016 IIBYT-CONICET. Octavio Giayetto is recipient postdoctoral scholarship from CONICET. Agustin Quaglia is Assistant Researcher at CONICET.

## Data, scripts, code, and supplementary information availability

All data and scripts supporting the findings of this study are available as supplementary material.

## Statement of authorship

GO, NFN and DA designed the research, GO, AEQ and MAP collected data, GO and AEQ performed the statistical analysis, all authors analysed the output data. GO wrote the first draft of the manuscript and all authors contributed with the manuscript revision.

