## Supplementary material SM1 for "Integrated environmental drivers and evolutionary history reveal conserved patterns of avian exposure to West Nile and Saint Louis encephalitis viruses"

**Supplementary Material 1.**

Ecological and life-history traits data.

Pace of life strategy.

The first axis of a principal component analysis (PCA) was used to depict the position of species along the spectrum of slow-to-fast life-history variation performed on nine variables on 9 variables related to reproduction (egg mass, clutch size and broods per year), size (wing length, tarsus length and body mass), and development (incubation period, generational length and fledgling period). The first axis (PCA 1) explained 54.8% of the variability and represented a gradient from fast (negative values) to slow (positive values) life-history strategies (Supplementary Fig. SM 1.1). Loadings suggest that wing length (91 %), egg mass (90%) and body mass (84%) contributed the most to the PCA 1 axis.

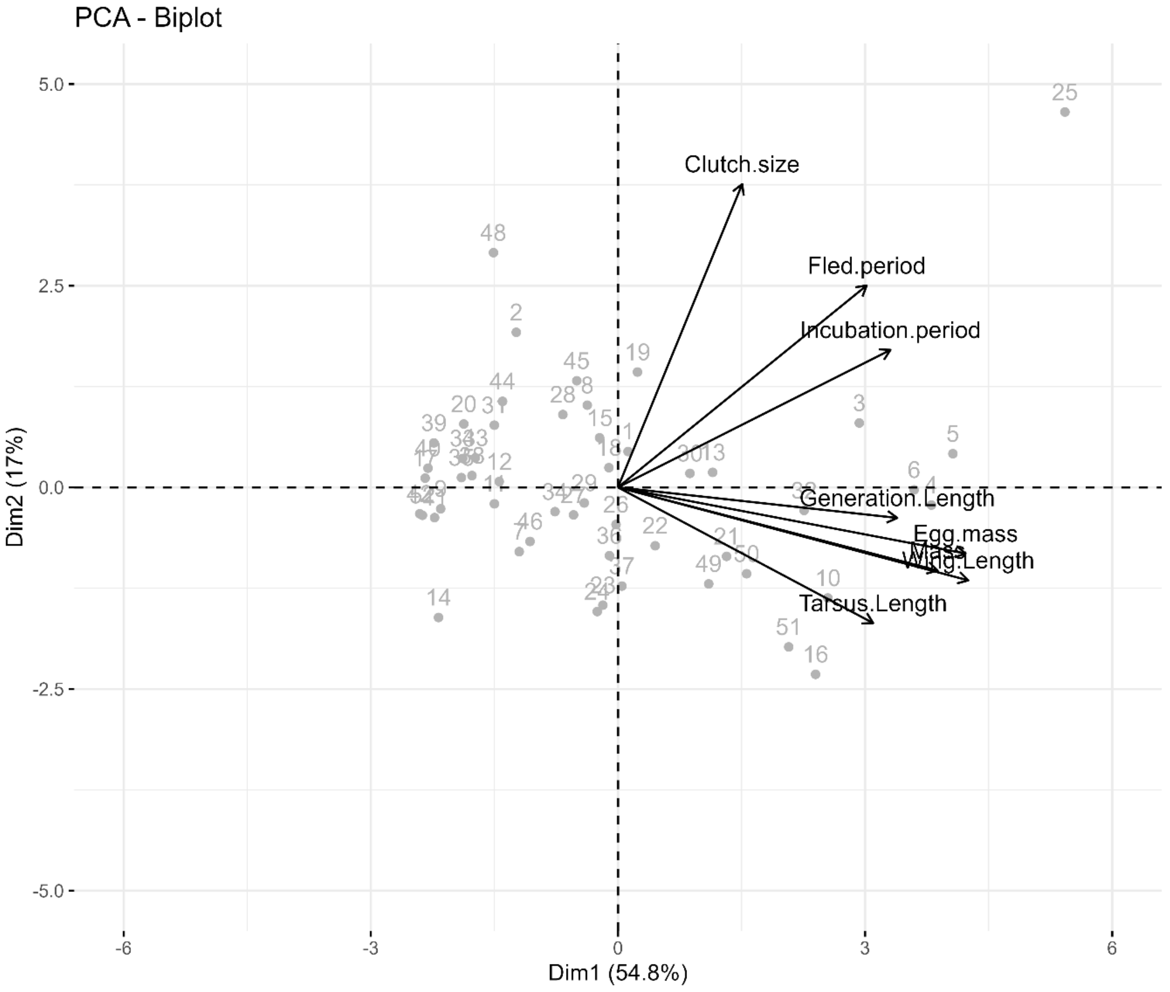


**Supplementary Fig. SM 1.1.** Principal Component Analysis (PCA) plot illustrating ecological traits of birds' species. Dots represent bird species in the spectrum of slow-to-fast life-history strategies and vectors represent the distribution of the ecological traits.

Spatiotemporal climatic variability in sampling sites.

To account for the variability in each sampling event, specific bioclimatic data from each site, season, and year were used, resulting in a total of 31 independent sampling events. We used environmental information that could affect arboviral transmission network to characterize the environmental niche where viral transmission occurs (Giesen et al. 2023). Hereafter referred to as Environmental Scenarios of Viral Transmission (ESVT). We extracted precipitation, minimum and maximum temperatures for each sampled month, and annual mean temperature, temperature seasonality, annual precipitation, annual water vapour pressure, annual wind speed for each sampled year and meters above sea level for each sampled site from https://www.worldclim.org/. We included the landscape characteristic for each sampled site following the next categories: Urban, agroecosystem, bosque chaqueño, chaco árido, chaco serrano and wetland.

The 31 sampling events were ordered according to their bioclimatic characteristics by means of a principal coordinate analysis (PCoA) using Gower distance matrix (Supplementry Fig. SM 1.2). From the PCoA results, we kept the number of axes that retained at least an 80% of the bioclimatic variability (Supplementary Fig. SM 1.3.A). After obtaining a new coordinate matrix with 8 axes (corresponding to the 80% of bioclimatic variability), we performed a two-stage k-means clustering analysis. We initially set the number of clusters and the starting points and then used this result to establish the optimal number of groups a posteriori, using the number of clusters that relatively minimize the intra-cluster variation (Supplementary Fig. SM 1.3.C). Finally, we obtained 7 clusters of sampling events grouped by their environmental resemblance (Supplementary Fig. SM 1.3.B).

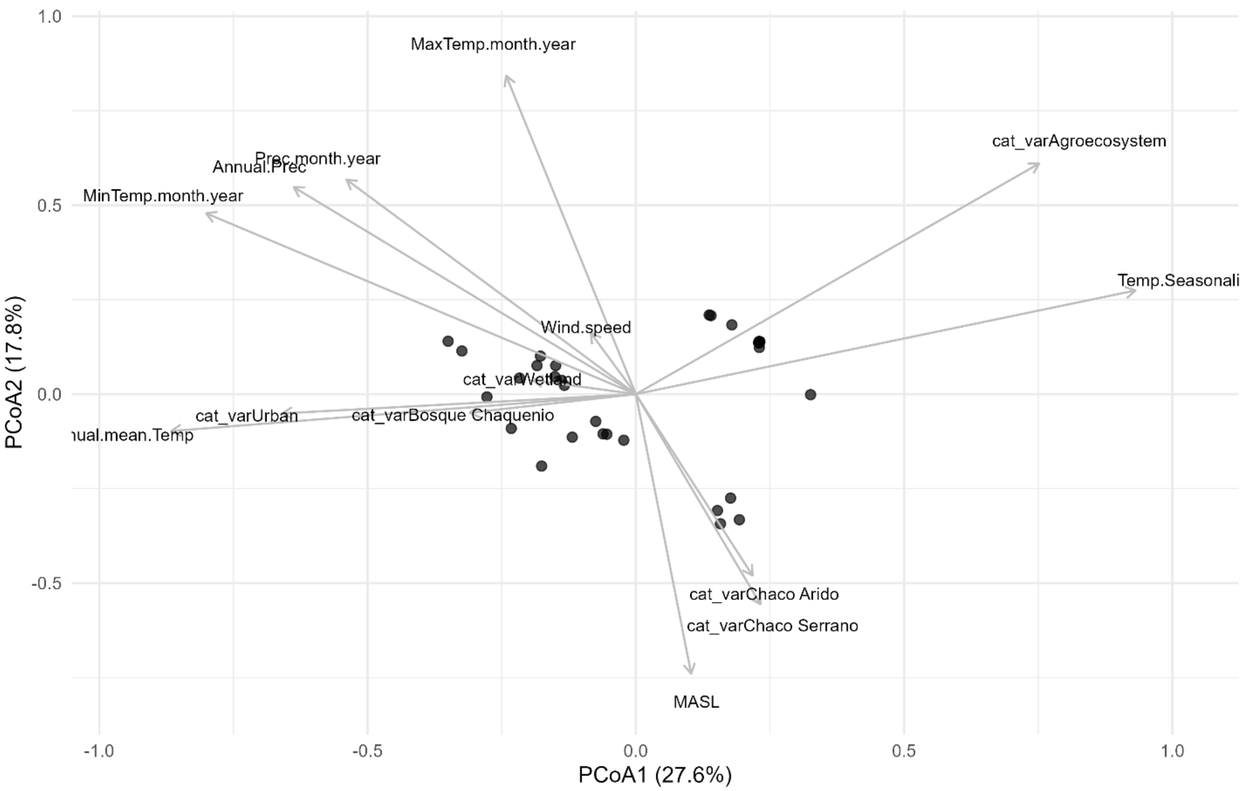


**Supplementary Fig. SM 1.2.** Principal Coordinate Analysis (PCoA) plot illustrating the Environmental Scenarios of Viral Transmission (ESVT). Dots represent the 31 independent sampling events, and their position reflects environmental dissimilarities based on bioclimatic and landscape variables.

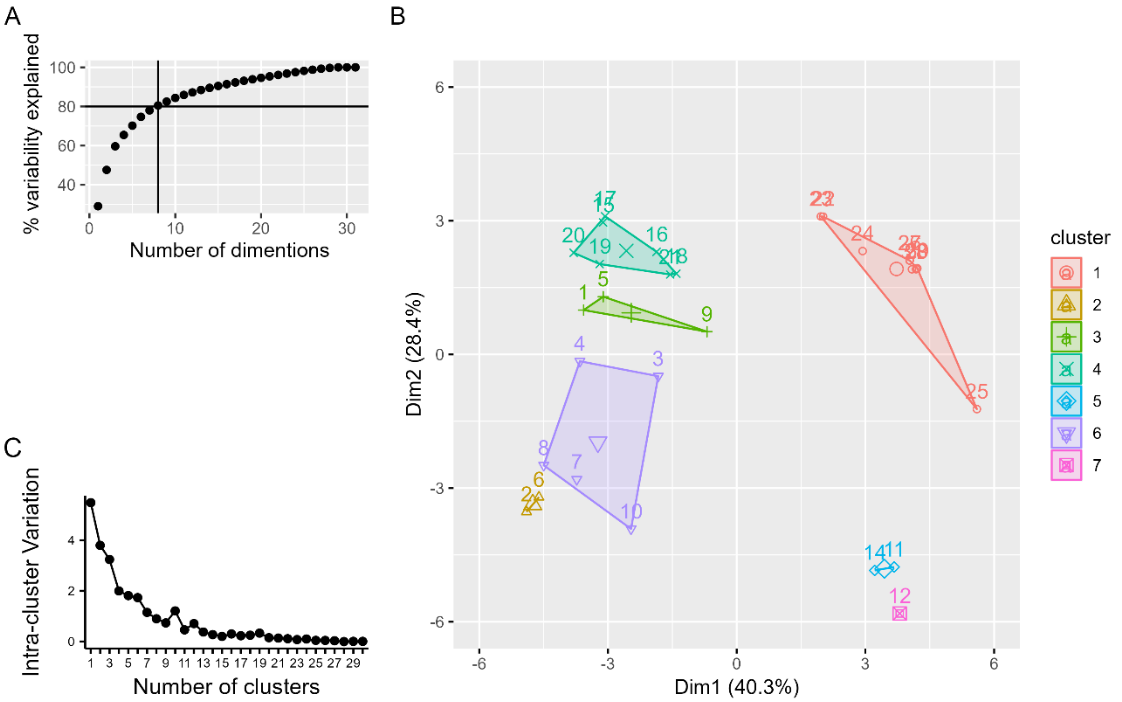


**Supplementary Fig. SM 1.3.** A) Number of axes that retained at least an 80% of the environmental variability. B) K-means clustering analysis of the selected number of dimensions in panel A and the selected number of environmental scenarios of viral transmissions that minimize the intra-cluster variation in panel C. C) Elbow method to obtain the optimal number of clusters that relatively minimize the total intra-cluster variation.
