## Supplementary material SM2 for "Integrated environmental drivers and evolutionary history reveal conserved patterns of avian exposure to West Nile and Saint Louis encephalitis viruses"

**Supplementary Material 2.**

Distribution of bird species, families and traits along environmental scenarios of viral transmission

The sampled coverage of the bird assemblage mirrored biodiversity organization along the biogeographic gradient analyzed. The number of species sampled varied consistently with the latitudinal gradient and the complexity of the ecoregions considered ($\chi^2 = 58.37$, df = 6, $p < 0.001$), whereas the number of families sampled was similar among ESTV ($\chi^2 = 9.64$, df = 6, $p = 0.140$) (Supplementary Fig. SM 2.1). The highest number of species was recorded in ESTV 4, being approximately twice that of ESTV 5 ($t = 4.14$, df = 23, $p = 0.008$) and three times that of ESTV 7 ($t = 4.26$, df = 23, $p = 0.006$) and 1 ($t = 6.17$, df = 23, $p < 0.001$). The distribution of ecological traits was consistent with expectations at the biogeographic scale analyzed (Supplementary Fig. SM2.2), as supported by the fourth-corner analysis indicating a random distribution of traits across ESTVs (Table. SM 2.1).

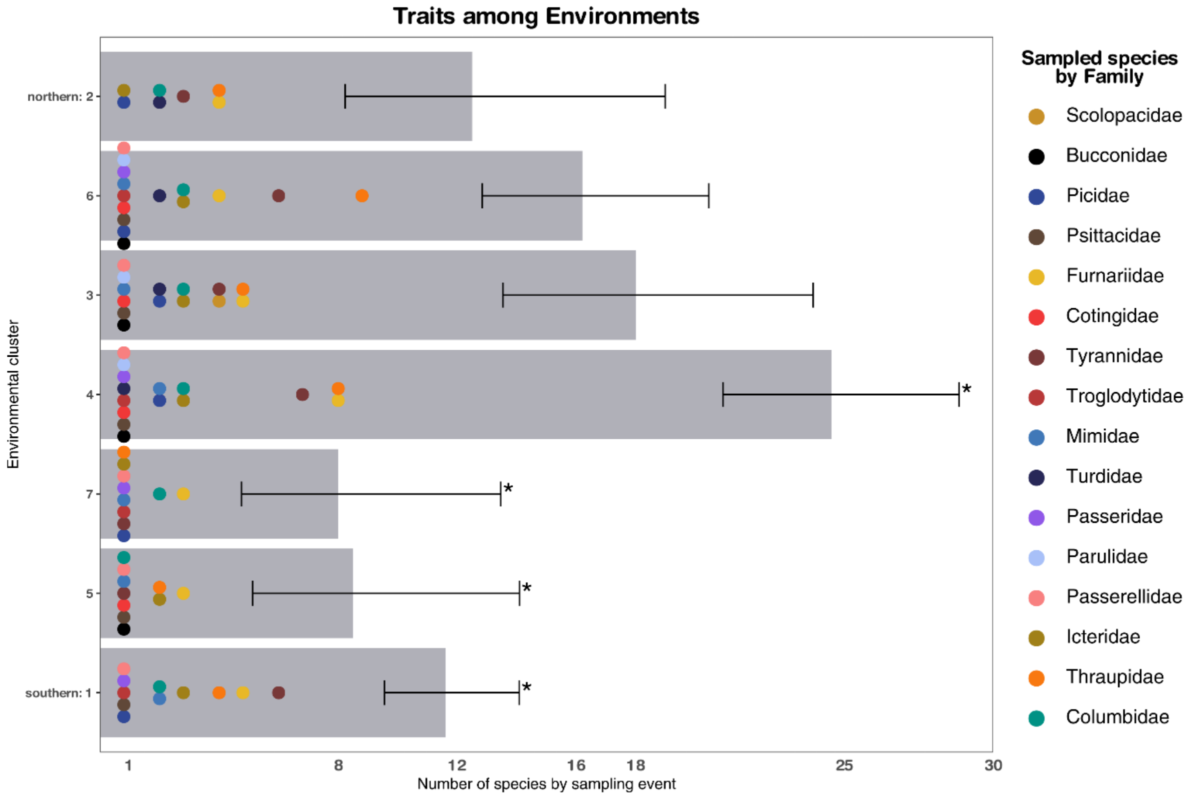


**Supplementary Fig. SM 2.1**. Distribution of bird species and families across the environmental scenarios of viral transmission (environmental clusters). * Significant post hoc multiple comparisons ($p<0.05$) after Bonferroni adjustment of the number of species sampled.

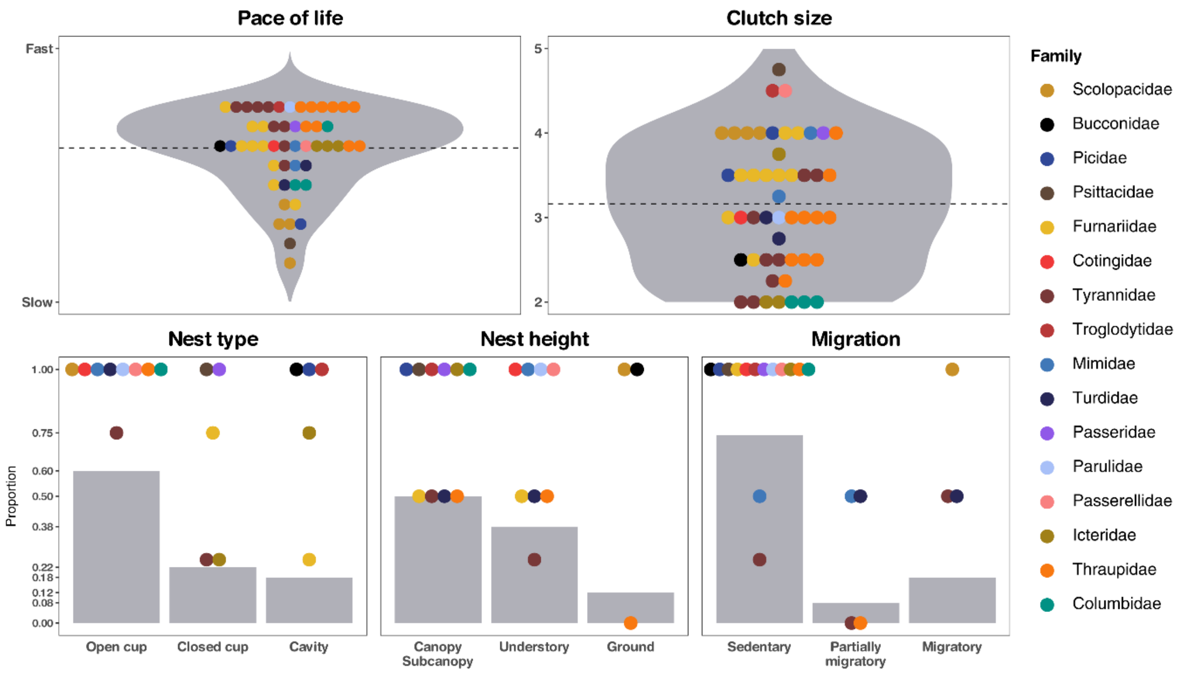


**Supplementary Fig. SM 2.2.** Distribution of sampled bird traits across families. Dashed lines indicate mean trait values across the assemblage.

**Supplementary Table. SM1.1**. Species trait–environment relationships under viral transmission scenarios. Fourth-corner analysis with Monte-Carlo permutation test.

| Species traits-environmental relationships | Statistic | Value (std) | Adjusted *p* |
|---|---|---|---|
| ESTV/ Pace of Life | pseudo-F | 51.55 (1.15) | 0.204 |
| ESTV / Clutch size | pseudo-F | 78.36 (2.62) | 0.061 |
| ESTV / Nest type | $X^2$ | 669.72 (2.55) | 0.061 |
| ESTV/ Nest height | $X^2$ | 298.00 (-0.18) | 0.515 |
| ESTV / Migration | $X^2$ | 306.13 (-0.04) | 0.515 |

Infection risk among Environmental Scenarios of Viral Transmission.

The posterior median infection risks (with 95% highest density intervals, HDI95%) for SLEV and WNV are shown across the seven Environmental Scenarios of Viral Transmission (ESVTs). For SLEV, the highest posterior mean was in Pampeano Occidental–Espinal (ESVT 1: 0.382 [0.159, 0.701]), followed by Chaqueño Oriental–Sabanas Inundables del Paraná (ESVT 2: 0.241 [0.073, 0.520]), Espinal (ESVT 3: 0.229 [0.089, 0.414]), Espinal–Chaqueño Occidental (ESVT 6: 0.218 [0.083, 0.426]), Chaqueño Serrano (ESVT 7: 0.162 [0.045, 0.410]), Espinal (ESVT 4: 0.166 [0.069, 0.339]), and Chaqueño Occidental (ESVT 5: 0.137 [0.036, 0.338]). For WNV, the highest posterior mean was in Chaqueño Oriental–Sabanas Inundables del Paraná (ESVT 2: 0.226 [0.029, 0.738]), followed by Espinal (ESVT 3: 0.217 [0.043, 0.704]), Pampeano Occidental–Espinal (ESVT 1: 0.213 [0.031, 0.712]), Chaqueño Serrano (ESVT 7: 0.164 [0.022, 0.674]), Chaqueño Occidental (ESVT 5: 0.165 [0.021, 0.631]), Espinal (ESVT 4: 0.103 [0.016, 0.471]), and Espinal–Chaqueño Occidental (ESVT 6: 0.090 [0.013, 0.439]) (Supplementary Fig. SM 2.3).

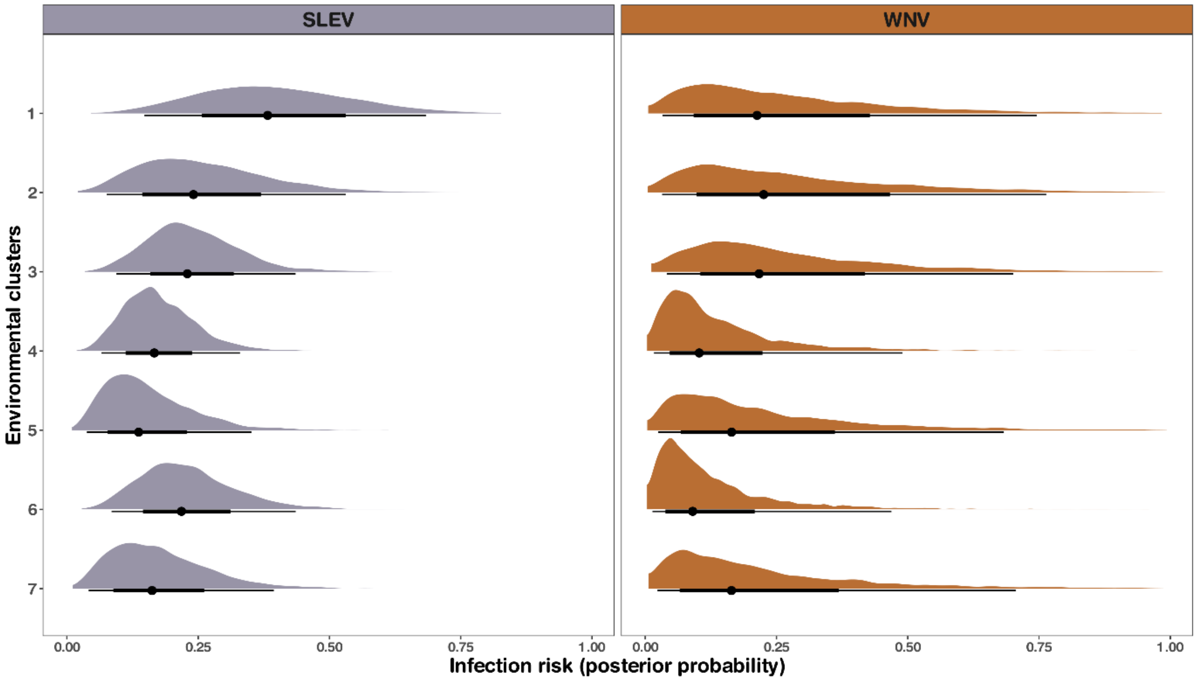


**Supplementary Fig. SM 2.3.** Posterior median infection risks for SLEV and WNV across the seven Environmental Scenarios of Viral Transmission (ESVTs). Each point represents the posterior median, and the lines indicate the 50%, 75%, and 95% highest density intervals (HDI).
